# Virtual reality as a tool for environmental conservation and fundraising

**DOI:** 10.1101/785014

**Authors:** Katherine M. Nelson, Eva Anggraini, Achim Schlüter

## Abstract

Anecdotal evidence from philanthropic fundraisers shows that virtual reality (VR) technology increases empathy and can influence people toward pro-environmental behavior. Non-profit organizations are increasingly marketing their causes using virtual reality and they report increased donations when VR technology is employed. In VR, users are immersed in situations intended to feel more like the real world through technology, such as 360° video screened through 3D headsets that block out visual and auditory distractions. The framing of the message as either positive or negative has long shown to have an effect on behavior, although consensus on the impact of framing has not been reached in relation to encouraging contributions to public goods. This paper focuses on field experiments used to investigate the effects of varying degrees of visual immersion and positive versus negative message framing on respondents’ contributions to a conservation charity. Participants were exposed to a five-minute underwater film about coral reefs and the importance of protecting them. We employed a 2×2 experimental design using 3D head-mounted displays comparing 360° film footage vs. unidirectional film and a positive message vs. a negative message. After watching the film, each participant completed a short questionnaire and had the opportunity to donate to a marine conservation charity. In addition, we tested a control treatment where no video was observed. The video was filmed in Indonesia which is host to some of the world’s most biodiverse reefs that are under great threat from human activity. We also conducted the study in Indonesia, sampling a total of 1006 participants from the Bogor city area and tourists on the island of Gili Trawangan - which is popular for scuba diving and snorkeling. We find significant differences in observed behavior and reported emotions between all treatments compared to the control condition. Among the tourist sample, we find significant differences between the 360° film with a negative message which garnered significantly larger average donation amounts compared to the unidirectional film with both positive and negative framing. Overall, we can infer from these studies that virtual reality is an effective way to raise awareness of environmental threats and encourage behavioral action, especially when tailored to target groups. New technology, such as the VR head-mounted display, is highly effective at attracting interest which is an important point to encourage organizations to invest in new technologies.

## 1. Introduction

The Intergovernmental Panel on Climate Change (IPCC) recently issued its special report on the impacts of global climate change on nature and society (Masson-Delmotte et al. 2018). The report paints a grim picture of the consequences of climate change if the earth’s temperature rises by even 0.5°C and further states that rising temperatures will result in food shortages, more wildfires, and a mass die-off of coral reefs as soon as 2040. Despite growing attention in the media pointing to the human-induced rate at which the climate is changing, mounting evidence from across the behavioral sciences has found that most people regard climate change as a non-urgent and psychologically distant risk—spatially, temporally, and socially—which has led to deferred action and decision making about mitigation and adaptation responses (van der Linden, Maibach, and Leiserowitz 2015).

Among the greatest challenges of communicating climate science to the public is bridging the knowledge-to-action gap (Moser and Dilling 2011). This gap refers to the general lack of environmental behavior change by individuals or society at large despite increases in communication about environmental problems and heightened public awareness of such issues. According to several studies, communication methods that are successful in bridging this gap involve cognitive and behavioral dimensions (Lorenzoni, Nicholson-Cole, and Whitmarsh 2007, Moser and Dilling 2011; Nelson et al. 2018a, 2018b). For example, people gain an understanding about an issue, experience an emotional response, such as interest or worry, and actively respond by changing climate-relevant behavior or political action. Research has shown that the content of the message (Tversky and Kahneman 1975, Gifford and Comeau 2011), and level of visual immersion (Shin 2018, Innocenti 2017, Rosenberg, Baughman, and Bailenson 2013) can have an impact on the intensity of emotional responses and influence behavior. Recently the nature documentary series *Our Planet* has received a plethora of media attention for the use of the negative framing of human-induced climate change impacts on nature which the series uses to push viewers toward a conservation website “to discover what we need to do now to protect our jungles” (Lowry 2019). Beyond framing, virtual reality^1^ (VR) technology has received attention for being able to increase immersion and one’s sense of presence, which is thought to affect emotions and connect people better to the subject matter (Diemer et al. 2015).

In this study, we examine virtual reality using a field experiment with real financial decisions. Using a head-mounted display, participants experience the five-minute video, *Coral Reefs: Life Below the Surface*^2^ (Schmitt et al. 2017). Then participants are asked to fill out a short survey which includes a request for donations to a coral reef conservation organization. Aiming to test the effects of visual immersion (low vs. high) and message framing (positive vs. negative), participants are exposed to one of the treatment conditions (see Table 1 for descriptions): positive video (VID_POS), negative video (VID_NEG), positive 360° video (360_POS), negative 360° video (360_NEG), and no video (control).

**Table 1.** Treatment conditions showing 2×2 experimental design (positive vs. negative framing) × (low vs. high immersion)

| Treatment conditions and description |  | Audio message frame |  | Visual Immersion |  |
| --- | --- | --- | --- | --- | --- |
|  |  | Positive | Negative | Low | High |
| VID_POS | video filmed with one camera (unidirectional) with positive audio | x | - | x | - |
| VID_NEG | video filmed with one camera (unidirectional) with negative audio | - | x | x | - |
| 360_POS | 360° video filmed with multiple cameras (omnidirectional) with positive audio | x | - | - | x |
| 360_NEG | 360° video filmed with multiple cameras (omnidirectional) with negative audio | - | x | - | x |
| CONTROL | No video | - | - | - | - |

## 2. Literature Review

The research concept to experimentally test the impact of different levels of immersion (low vs. high immersive environments) on emotional response and pro-social behavior was born out of the recent hype in popular media articles that virtual reality “can make you a better person” and is “the ultimate empathy machine” (Samit 2017). Such articles make claims that virtual reality increases empathy and the technology is successful at influencing behavior, such as increasing donations for charity (e.g. The Wall Street Journal “Charities Use Virtual Reality to Draw in Donors” (West 2015); AR Post “Using Virtual Reality for Increased Charity Donor Outreach and Funding” (Chang 2018); and The Guardian “Can Virtual Reality Emerge as a Tool for Conservation?” (Millar 2016)). However, at the point of conceptualization of this research, only anecdotal evidence was being used to support these claims. There is currently a lack of scientific research that can back up claims that increasing the level of immersion through the use of virtual reality technology can influence subsequent behavior, such as increased donations to charity (as the articles claim it can).

### 2.1. Immersive visual experiences

An article by Fortune magazine in 2017 claimed, “the emotional impact of VR has proven to increase awareness, evoke empathy, and elicit action” pointing to examples from a Facebook report that “48% of virtual reality charity content viewers were likely to donate to the causes they experienced” and a report by the United Nations claiming that their VR production *Clouds over Sidra* which films the life of a 12 year old Syrian refugee helped raise twice the charity’s normal rate” (Samit 2017). Although these studies may sound promising, they lack controlled experimental designs and many uncontrolled factors could account for the supposed increase in donations (e.g. increased media attention of the Syrian crisis may have led to higher donations that year regardless of the VR technology used at the fundraiser). The study from the aforementioned Facebook report and one from the market research company Nielson (2017) found donation intentions to be higher following exposure to a 360° video compared to other media, but these considered hypothetical donation decisions only.

Several lab studies performed at the Virtual Human Interaction Lab of Stanford University focus on the effects of virtual reality on encouraging prosocial behavior (Rosenberg, Baughman, and Bailenson 2013) and environmental behavioral intentions (Ahn et al. 2016). The evidence seems promising that virtual reality interactions increase a person’s feeling of presence which can connect them better to the subject matter and affect one’s behavior. In one study, Ahn, Bailenson, and Park (2014) compare the effects of cutting a tree in virtual reality to reading a written description or watching a video depiction of the tree-cutting process to encourage paper conservation. The study shows that the virtual experience results in a 20% decrease in paper use compared to participants who read a print description of tree cutting^3^. In another experiment by Ahn et al. (2014), the participants wear a virtual reality headset and can walk around on all fours to experience what it’s like to be a cow that is raised for dairy and for meat. As the participant wanders around the virtual stall with hay, he can see the embodiment of himself as a cow get poked with a cattle-prod while he is also actually experiencing a slight poke in the ribs while he is being led to the slaughterhouse in the virtual world. For a time after the VR experience, participants reported eating less meat (Ahn et al. 2016). On the contrary, preliminary research carried out by the authors of this paper found no statistical differences among university students exposed to differing levels of immersion in an underwater virtual world in relation to their charitable giving behavior to coral reef conservation (Nelson 2019). While some scholars believe virtual reality can create empathy with non-human actors by simulating, for example, what it’s like to be a cow or a piece of coral (Ahn et al. 2016), other scholars are highly critical of this concept (Ramirez 2017).

A lab study by Gürerk and Kasulke (2018) tests 360° videos using VR headsets and finds that participants exposed to VR gave more frequently and donated higher amounts compared to those that received only a written donation request with no video. The research described herein differs from the study by Gürerk and Kasulke (2018) in several important ways. We test the effectiveness of 360° films on raising money for charity outside of the lab using a natural field experiment. We control for the presence of the head-mounted technology ensuring that any differences in our results are driven by the immersive experience provided by the 360° video compared to a non-360° video so we know it’s not the new technological gear driving the result. And finally, we compare the combined effects of the level of immersion with a positive or negative audio message about the future of coral reefs based on climate change predictions.

### 2.2. Framing: Positive versus negative message valence

Framing in this context is defined as a semantic restructuring of identical information (Hallahan 2008). Valence framing structures information in either a positive or a negative light. In this manuscript, positive message valence describes action that leads to a favorable outcome (e.g., save coral reefs and species biodiversity) and negative message valence indicates the consequences of inaction which lead to adverse conditions (e.g., loss of coral reefs and species biodiversity). Classic economic theory of rational choice assumes that as long as the same information is presented, regardless of how the information is presented or framed, an individual will always interpret the information in the same way and behave accordingly (Avineri and Waygood 2013). Thus, preferences should not be affected by framing. However, a multitude of studies have proven otherwise, beginning with the seminal work of Tversky and Kahneman (1975), which eventually led to a Nobel Prize in economics in 2002, and which describes a cognitive bias known as “the framing effect” wherein people’s decisions are effected based on positive or negative semantics (i.e., people make different decisions if the options are presented as gains or losses).

Communicating climate change to encourage environmentally friendly behavior is crucial in transforming knowledge into action. The way climate change messages are framed can significantly alter the impact they have on a recipient, but not much is known about the effects of positive versus negative message valence on environmental attitudes and even less is known about the effect they have on actual behavior (Spence and Pidgeon 2010). There is a surprising scarcity of empirical research examining the effects of message framing on pro-environmental behavior (Cheng, Woon, and Lynes 2011). Many studies examining the effects of framing have been conducted in the field of health psychology which focus on comparing the relative effectiveness of information that focuses on the positive consequences of performing a particular behavior (gain) or on the negative consequences of inaction (loss) (Gong et al. (2013). However, a review of the literature on positive and negative message framing proves to be rather inconsistent and highly contextual with some studies showing that positive messages are more effective and other studies showing increased effectiveness with negative messages.

According to a health study by Rothman et al. (2006), evidence indicates that negative loss frames are found to be more effective for encouraging detection behavior and positive gain frames are better for encouraging prevention behavior. There is less empirical evidence, however, when communicating climate change risks as to whether insights and theories from other domains are transferable to climate action (Pelletier and Sharp 2008). Additionally, some studies suggest that framing effectiveness differs based on a person’s decision stage (i.e., early stage of determining whether an issue is problematic based on its personal risk/costs or the later stage of establishing intention to act). When personal risk to an individual has yet to be well-established, fear-based messages seem to be more effective at eliciting emotional responses (Pelletier and Sharp 2008, Rothman et al. 2006); whereas, once a person shifts from considering risks associated with a potential future situation to considering the potential solutions to avoid said scenarios, positive messages that emphasize a desired outcome and the benefits of adopting a specific behavior are more salient because these messages are now congruent with the actions that could eliminate risk or the fear associated with a specific issue (Hardisty and Weber 2009, O’Keefe and Jensen 2007). The inconsistency across studies supporting, in some cases, a stronger influence using negative loss frames and, in other cases, a stronger influence with positive gain frames, seems to point to additional influencing factors. According to a review of the literature, these include: the purpose of the information campaign (i.e., raise awareness, change opinions, invoke emotion, call-to-action to avoid or to do something, plan for future behavior); stage of individual decision process (i.e., risk consideration, prevention, detection, goal setting, short-term action, long-term behavior change); the particular behavior in question (i.e., actively doing something or avoiding a particular behavior, personal action, political action, societal action); the relationship between an individual and the behavior in question (i.e., demographics, personality type) (Spence and Pidgeon 2010, Pelletier and Sharp 2008, O’Keefe and Jensen 2007).

Most research of valence framing does not consider social dilemmas such as climate change, and the nature of impacts of framing may be different in such situations (Avineri and Waygood 2013). A theoretical study by Moser and Dilling (2007) proposes shifting the discourse towards a positive motivational-oriented approach offering solutions rather than the negative approach that highlights sacrifices as a more effective strategy to encourage climate-change-related behaviors. Gifford and Comeau (2011) empirically test this using a motivational (positive) or sacrificial (negative) priming message in a survey eliciting intentions to engage in mitigative behaviors and they find that the positive message is more effective at encouraging several environmental behaviors compared to the negative message, but these results are not significantly different than the control condition. Several demographic factors such as gender, age, income, and education level also play a role in determining behavior. A study by Avineri and Waygood (2013) examines the effect of positive and negative valence framing on people’s perceptions of CO2 emissions based on how a travel mode is framed (i.e., it provides environmental benefit (positive frame) or its potential to reduce environmental loss (negative frame). The study finds that negative framing is more effective at highlighting the differences between CO2 emission amounts of alternative travel modes which they suggest is more likely to influence travel-related decisions. These studies are highly informative and begin to fill the gap in the literature on climate-change-related behavior, although more research is need that pushes beyond hypothetical responses and intended behavior. Given the dearth of empirical evidence in this field, the research presented in this paper measuring the effects of framing on emotions, intentions, and actual pro-environmental behavior is timely and fills a crucial gap in research.

## 3. Methodology

### 1. Study location

Indonesia is the largest archipelagic country in the world, encompassing more than 17,000 islands, 86,700 square kilometers of coral reefs, and supporting nearly 230 million people. The people of Indonesia are increasingly dependent on marine resources for food and income sources, including tourism. Presently about 70% of the country’s protein sources come from fish (in some poor coastal communities this figure approaches 90%). In 2016, total contributions from tourism in Indonesia amounted to 60 billion USD in revenue (6.2% of GDP) (BPS 2018). Indonesia is a global hotspot for coastal marine biodiversity. Blessed with nearly 18% of the world’s coral reefs, Indonesia sits firmly at the center of the “Coral Triangle”, the region with the world’s highest marine biodiversity (Hughes, Bellwood, and Connolly 2002). Despite the clear importance of coral reefs, they are rapidly declining as a result of a range of local drivers (e.g., overexploitation of fisheries, coastal development and pollution) and global drivers (e.g., global warming and ocean acidification) (Hoegh-Guldberg 2011). Conservation of coral reefs and the resources they provide is a collective responsibility and requires action from people in countries across the globe to reduce greenhouse gas emissions. Locally, efforts to reduce additional pressure from already stressed reefs can improve their resilience to climate change (Hughes et al. 2003).

The study was conducted in two locations in Indonesia. We sampled students from Bogor Agricultural University (Institut Pertanian Bogor - IPB), a general population from a public park in Bogor, and a mostly foreign tourist sample that were visiting the popular scuba diving destination of Gili Trawangan (see Figure 1). We chose these locations and sample populations with our research collaboration partners in Indonesia (IPB and Gili Eco Trust) to cover 1) the typical lab experiment student population, 2) a more generalized public population, and 3) a targeted population.

**Figure 1.**
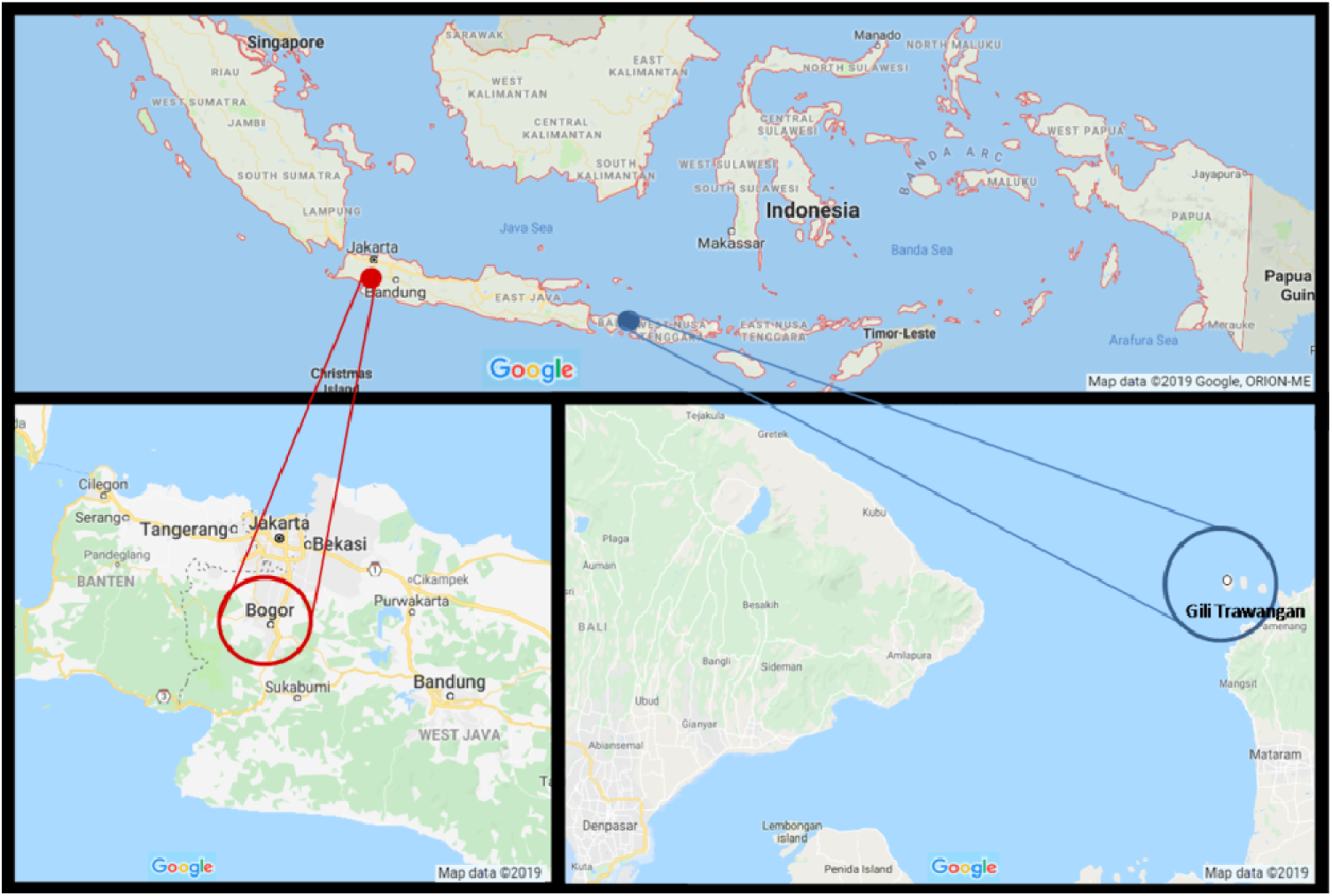
Map highlighting (top) Indonesia, (left) Bogor, and (right) Gili Trawangan. Source: Google maps 2019.

Bogor has a population of approximately one million people and is located 60km from the capital city Jakarta on the island of Java. Bogor is an important economic, cultural, and scientific center for Indonesia. Bogor is a land-locked city and is situated in the mountains, at a distance, relatively-speaking, from the popular coral reef diving destinations of Indonesia. In contrast, Gili Trawangan, a small island situated between Bali and Lombok, is represented by a remarkably biodiverse coral ecosystem. Over the last decade Gili Trawangan has grown into a major destination for tourists worldwide of all budgets and interests (Nelson et al. 2019). Gili Trawangan is the second most frequented destination in South East Asia for SCUBA diving certification (second only to Koh Tao in Thailand) (Partelow and Nelson 2018). Although only six square miles, the island receives heavy tourist traffic with up to 2000 new visitors per day and over one million tourists annually (Halim 2017). Increased tourism results in rapid development, challenging the island’s infrastructure to keep up with the growth and putting pressure on the fragile coral ecosystem surrounding the island (Partelow and Nelson 2018).

Field experiments were conducted during March and April, 2018. All participants watched a five minute video and then completed a short questionnaire, except for those in the control condition that only completed the survey. The experiment employs a between-subjects 2×2 design examining the effectiveness of increased visual immersion through 360° film technology compared to classic unidirectional film footage and positive or negative valence framing (see Table 1 for treatment descriptions). We measured differences in donation behavior and emotions post-treatment across the four treatments and the control (no video).

In total, 1006 subjects participated in the experiment. 487 subjects were recruited from the IPB campus and 248 people from the general public were recruited from an outdoor public park in the center of Bogor. We then conducted the experiment with 271 participants on the popular diving and tourism island Gili Trawangan. See Table 2 below for details on sample size by treatment and location.

**Table 2.** Treatment group and sample size by location.

|  | **VID_POS** | **VID_NEG** | **360_POS** | **360_NEG** | **CONTROL** | **TOTAL** |
| --- | --- | --- | --- | --- | --- | --- |
| Student - Bogor | 93 | 98 | 97 | 99 | 100 | **487** |
| Public - Bogor | 49 | 51 | 50 | 48 | 50 | **248** |
| Gili Trawangan | 54 | 51 | 56 | 59 | 51 | **271** |
| **TOTAL** | **196** | **201** | **203** | **206** | **201** | **1006** |

Several field assistants were trained on the protocol and assisted with recruitment, video screening, and survey administration. Participants were randomly assigned into treatment groups based on random allocation of surveys that were secretly coded with an ID number which specified the treatment condition. The assistant then helped the respondent with the headset and headphones and started the video in the preferred language (the video was available in Bahasa, English, and German). The video lasted five minutes. After the participant watched the video, they filled out the survey questionnaire which included: a request for a donation to charity, questions about emotional feelings, environmental engagement, and basic demographics.

Respondents from Bogor were incentivized through enrollment in a lottery with a ten percent chance to win 100,000 Indonesian Rupiah (IDR)^4^ (approximately $7USD or 6€EUR). Participants were asked if they were to win the 100,000IDR, how much (if any) would they donate to the coral reef charity, Gili Eco Trust. For notifying the respondents in Bogor about the lottery winnings, the field assistant would text the ID code to the head researcher to check if the respondent’s number matched the randomly pre-selected winning ID codes. In the cases that the respondent won the lottery, the donation amount was deducted from their winnings. Lotteries, including raffles, are commonly accepted methods used in experimental economic research and in charity fundraisers (Carpenter and Matthews 2017). Respondents from Gili Trawangan were not offered the lottery incentive and were not paid for participation. The surveys in Gili Trawangan included an empty envelope with the corresponding ID number where the respondent would put their donation and drop the envelope into a collection box. The purpose of different experimental designs was to create plausible scenarios for each context and to reach sufficient donations in the Bogor sample, to possibly observe a treatment effect. Both designs were pretested.

The authors recognize that people may treat windfall earnings or possible future earnings differently than money in their own pockets. Therefore, we do not compare donation results across the Bogor sample and the Gili Trawangan sample. The donation data are treated and analyzed as two separate studies. We are interested in treatment differences and trends that may emerge across controlled conditions where all respondents were treated the same. To avoid overstating donations in the Bogor case and to make the results broadly applicable, we focus on significant differences in donations between treatments compared to the control condition rather than the actual donation amounts in the respective studies.

In all treatment conditions, we used the Zeiss VR One headset, Nokia 6 smartphone, and Sony MDRZX110 headphones to present the film to participants. We were particularly interested in understanding if the increased sense of immersion provided by 360° film technology influences charitable donation decisions compared to standard unidirectional film. 360° film viewed using a VR headset provides an immersive experience by surrounding one’s field of vision with continuous video imagery. Other studies have compared watching a 360° film on a mobile phone or a tablet to watching the film using a headset (Gürerk and Kasulke 2018). However, we want to know the effect of the increased visual immersion provided by the 360° omnidirectional film technology. In the study by Gürerk and Kasulke (2018), the new technology of the headset and the additional piece of equipment which completely obstructs ones vision thereby blocking out any other visual distractions may be enough to drive the differences they observe in donations between treatments. It is not possible to disentangle the effects of the equipment itself versus the level of immersion created by the full field of vision compared to a limited field of vision without testing both the classic unidirectional video and the 360° video with the headset on. Therefore, all participants, except those in the control condition, viewed the video wearing the VR headset.

### 3.1 Unidirectional vs. 360° film

Unidirectional film is shot with one video camera facing in only one direction at a time. The viewer has a limited field of view given that they are only able to see in one direction. The viewer is a passive participant in the visual story dictated by the filmmaker. Participants that were exposed to the unidirectional film using the VR headset would see the video projected onto a screen in front of them similar to watching a movie in a theater or on TV. The screen is rectangular and framed on a solid indigo background (Figure 2a). The participant could move their head in any direction but they would only see the indigo background in every direction except forward facing where the video is playing. 360° film is shot using an omnidirectional camera where a view in every direction is recorded simultaneously. The viewer has the freedom to look around in every direction throughout the video. When a 360° video is viewed using a VR headset, the movement of one’s head changes the direction of the view. This creates an immersive experience given that the video responds to the user’s movement. When the user looks right, they see video footage to the right, when they look down, they see the corresponding video footage showing the sea floor and when they look up, they see the surface of the water (Figure 2b shows a relative wider field of vision but cannot fully capture the 360° experience with a static two dimensional image).

**Figure 2.**
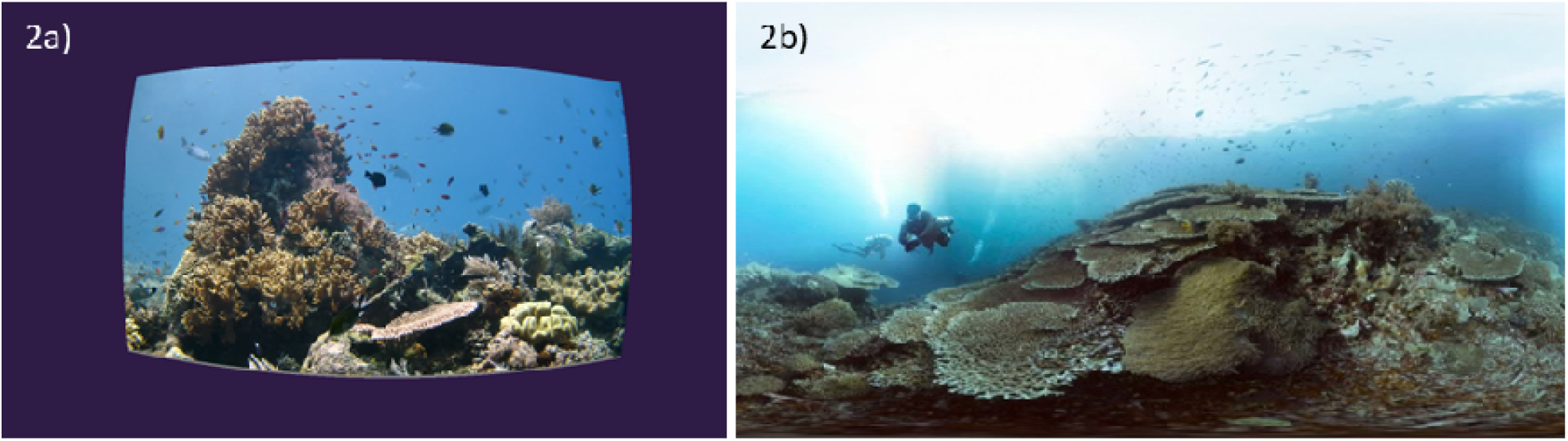
Screen capture of the field of view of unidirectional film (2a) and 360° film (2b) as seen while wearing the Zeiss VR One headset. Note: The 360° film imagery in 3b is a dynamic 360 degree effect which surrounds the user in every direction with video footage but this cannot be captured in a static image. Source: Schmitt, H. and Schmitt, C. [The Jetlagged]. *Coral Reefs: Life Below the Surface*. Retrieved from https://www.youtube.com/watch?v=2TPG8lcfeDc&t=64s

### 3.2 Positive vs. Negative message valence

To test the effectiveness of positive and negative valence, two scripts were semantically structured so that environmental gains and losses were distinctly presented. The positive valence narration focuses on the benefits that the reefs provide and will continue to provide in the future if we act now to save them. The negative valence focuses on the losses we will suffer if we fail to act (see Table 3 below for excerpts from the narrative script; see Appendix for full narrative). The imagery between the positive and negative valence framing was the same.

**Table 3.** Narrative script for positive and negative valence frame.

| Positive valence | Negative valence |
| --- | --- |
| Countless animal species, but also us humans depend on healthy coral reefs. | Countless animal species, but also us humans will suffer if coral reefs are destroyed. |
| They provide food for millions, security and protection from storm surges, sustain livelihoods and host the potential for new medicines. They are essential for the balance in the oceans. | Millions will lose their source of food and their livelihoods. Storm surges will hit the coastlines with full force. Potential cures for human diseases that might be found in the ocean could be lost forever. |
| We must protect it before it's too late. | If we don't protect the ocean now, it might already be too late. |
| Each person and each action makes a difference. | If we don't change the way we treat our planet, pollution and plastic waste will accumulate. Carbon dioxide will proliferate, driving global warming to higher and higher levels. |
| Together we can save the ocean for a bright and blue future of our planet. | If we don't act now, coral reefs will likely be gone within the next 30 years. |
| If we act now, we can save the coral reefs. |  |

### 1.1 Charitable organization receiving donations

Donations were collected for the coral conservation organization Gili Eco Trust. The Gili Eco Trust is a local community-organized conservation organization on Gili Trawangan that focuses efforts on restoring and regenerating coral reefs using Biorock technology (Goreau and Prong 2017). In addition to implementing and maintaining one of the largest coral restoration projects in the world, the Gili Eco Trust manages several sustainability programs on the island including weekly beach cleans, garbage and compost collection, plastic bottle bank (money for plastic), recycling services (sorting, cleaning, crushing, and shipping off the island to a recycle center in Lombok), sourcing alternative plastic products for island businesses, advocating for sustainable policies and infrastructure, and providing conservation education to the local schools. The Gili Eco Trust operates solely on donations from local businesses, tourists, and volunteers.

## 4. Results

The summary statistics of each sample population are described in Table 4 below. The mean age of participants in the study was 23 with the student sample representing a younger average age of 20 years old and the tourist population averaging 29 years old. On average there were slightly more female participants than males. The student sample came from locations that were closer to the sea with an average of 74km relative to the public sample at 200km and the tourist sample at 319km. Income was measured based on the participants’ perception of their family income in relation to the average income level from their country.

**Table 4.** Summary statistics of each sample and the overall sample.

| Demographic | Overall Total | Student - Bogor | Public - Bogor | Gili Trawangan |
| --- | --- | --- | --- | --- |
| *N* | 1006 | 487 | 248 | 271 |
| Age | 23.4 (8.26) | 20.4 (3.55) | 23.2 (9.22) | 29.3 (10.25) |
| Female (%) | 53 | 55 | 47 | 55 |
| KM to sea | 165 (700) | 74 (117) | 200 (1074) | 319 (880) |
| Income | 3.9 (1.3) | 3.8 (1.1) | 3.6 (1.6) | 4.2 (1.5) |
| Education | 4.4 (1.1) | 4.6 (0.9) | 3.6 (1.1) | 4.6 (1.1) |
| Religiosity | 4.7 (1.8) | 5.3 (1.1) | 5.4 (1.4) | 2.8 (2.0) |

Note: The numbers indicate means (standard deviations in parentheses) except where percentage (%) is indicated. Variables: *N* (sample size), Age (in years), Female (% of women), KM to sea (distance from one's home to the sea in kilometers), Income (perceived position in the income scale ranging from 1 being the lowest income to 7 being the highest income), Education (1 elementary school, 2 secondary school, 3 high school, 4 vocational training, 5 bachelor's degree and 6 post-graduate degree), and Religiosity (self-identified on a scale from 1 being not at all religious to 7 being the most religious).

Most people across the sample consider themselves to be in the middle income category with the general public sample perceiving themselves on average to be on the low end of the middle income category. The average level of education was the same for the student population and the tourist population, while the general public in Bogor were lower on average by one level of education. The level of religiosity was measured based on self-identification on a 7 point Likert scale of ‘not at all religious’ to ‘the most religious’. Participants from Bogor claim to be much more religious compared to the tourists from Gili Trawangan who overall self-identified as not very religious.

### 1.2 Bogor, Indonesia

The results of treatment differences based on the average contribution are reported for the student population from IPB and the general public from Bogor in Figure 3. There are no significant differences in the average contribution amounts between the two populations; therefore, the data are combined for the regression analyses.

**Figure 3.**
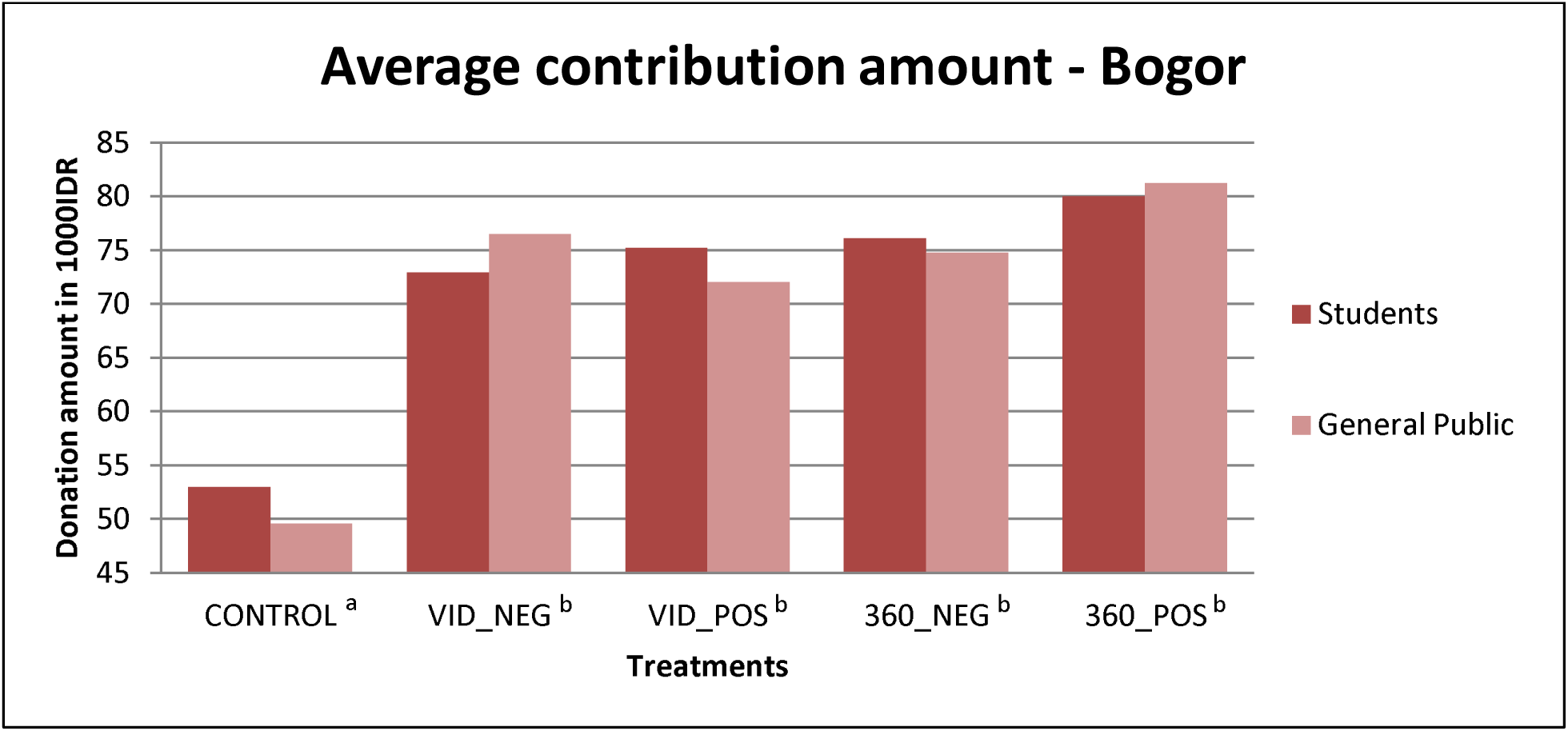
Average contribution amounts from students and the general public from Bogor. Treatment conditions with different superscript letters are significantly different at p<0.01; those with the same superscript letters are not statistically different from each other using the one-way ANOVA with Bonferroni correction for multiple comparisons.

Analyzing the results using the one-way ANOVA with Bonferroni correction for multiple comparisons, we find that average contribution amounts are significantly higher in all treatment conditions at p<0.001 when compared to the control condition. The four treatment conditions, however, are not significantly different from each other (see Figure 3). As can be seen from the bar graph, respondents in both the student sample and the general population from Bogor donated more on average in the 360° positive message frame treatment than the average contributions in the other treatments, however, this amount was not significantly higher when compared to the 360° negative treatment or to the classic unidirectional video treatments.

Upon examining donations on the extensive margin (i.e., to give or not to give), we must consider that donations would come out of any lottery winnings. Therefore, the frequency of donation was very high (>90%) in all conditions, including the control, with no significant differences between treatments. As reported in Figure 3, there is a treatment effect on the intensive margin (i.e., average amount given) but only when the treatments are compared to the control. We do not find a significant effect between positive and negative framing (VID_NEG vs. VID_POS and 360_NEG vs. 360_POS) or between high and low visual immersion (VID_NEG vs. 360_NEG and VID_POS vs. 360_POS)

For the linear OLS regression, we combined the data from the student population and the general population in Bogor since there are no statistical differences in their donation behavior. An environmental engagement index (Cronbach’s α = 0.68) was created from the combined mean score of all the environmental engagement indicators including the questions: *I prefer buying new things rather than fixing broken things* (reverse coded), *I prefer to eat meals with meat/fish rather than vegetable only* (reverse coded), *I bring my own reusable bag to the market, I recycle, I bring my own container for take out, I volunteer time for an environmental group, I use single use plastic* (reverse coded), *I make travel plans based on the amount of CO2 emissions generated by transportation, I purchase recycled materials, I only buy products from companies that protect the environment*.

The average donation amount is used as the dependent variable. Several iterations of the regression were analyzed by first including the demographic variables, then adding the environmental engagement and attitude variables, and adding the treatment assignments. The results show that age and income are significant indicators of donation amount and as age and income increase, donation amounts also increase. The behavioral intention indicator ‘sustainable future’ which asked participants to rank where they see themselves in the future in respect to living a sustainable lifestyle was also significantly positively correlated to the amount donated. However, after controlling for the treatment effects, the significance of the behavioral intention is lessened. Compared to the left out control treatment, all other treatments are highly significant determinants of the donation amount. The treatment variables are strong explanatory variables as can be seen by the increase in R^2^ when the treatments are added to the regression model. Other factors, such as gender; education; distance a person lives to the sea; perceived level of personal religiosity / climate-change optimism / generosity / environmental consciousness; and previous experience seeing a real reef do not seem to be significant drivers of donation decisions.

### 3.3 Gili Trawangan

The results presented in Figure 4 show the treatment differences on the extensive margin. The bar graph shows that the frequency of donations is higher in the virtual reality treatments. The rate of donating is significantly higher (p<0.001) in all treatment conditions when compared to the control condition using a binomial probability test. There are no significant differences between positive and negative framing in either the low visual immersion (VID_NEG vs. VID_POS) or high visual immersion (360_NEG vs. 360_POS) treatments. When comparing the effects of level of visual immersion, we find the rate of giving to be significantly higher (p<0.01) in the high visual immersion treatments for both the positive (VID_POS vs. 360_POS) and negative frames (VID_NEG vs. 360_NEG).

**Figure 4.**
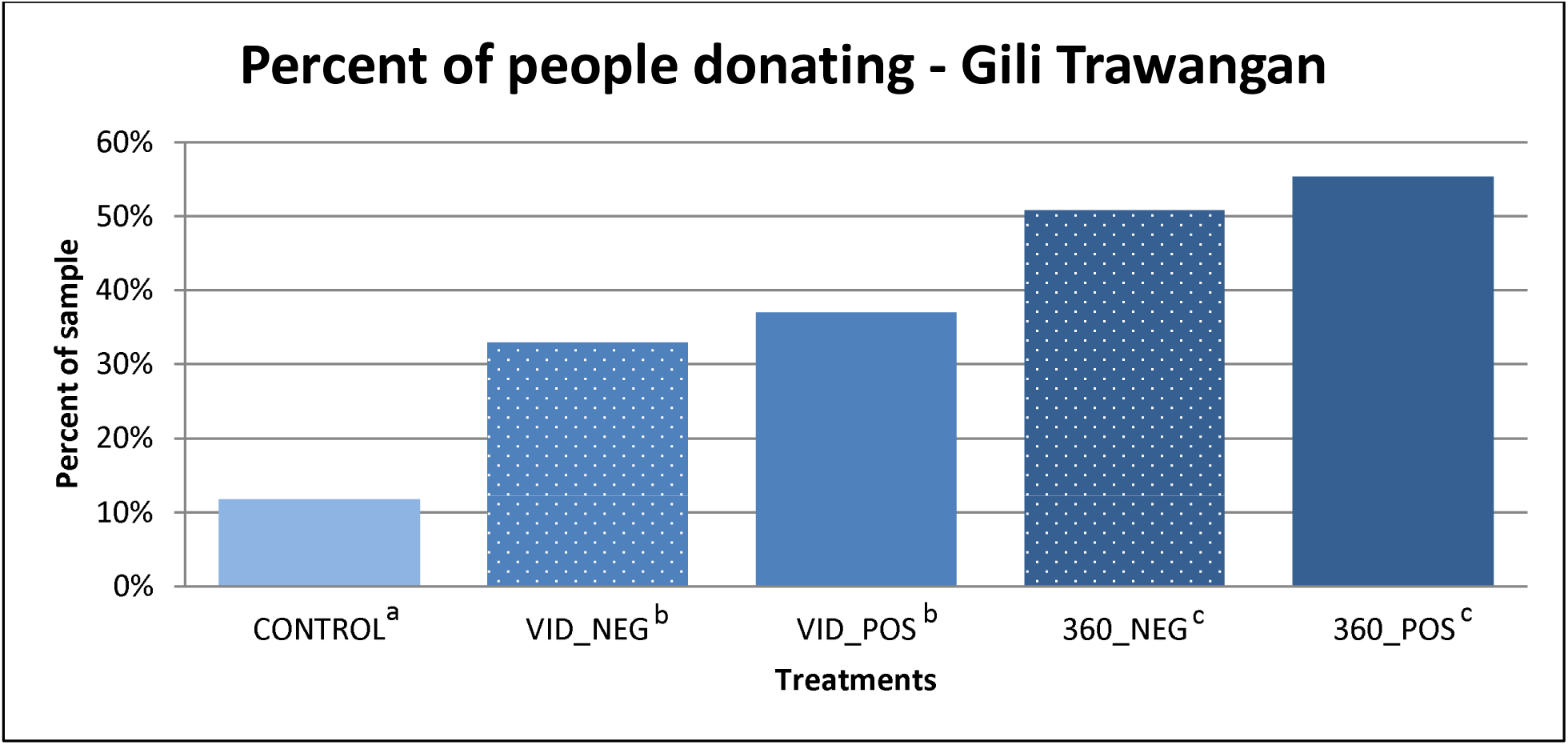
Percentage of participants that donate a positive amount to Gili Eco Trust. Treatment conditions with different superscript letters are significantly different; those with the same superscript letters are not statistically different from each other.

Figure 5 shows the results of the average donation amounts across the treatment conditions. There are significant differences (p<0.05) between both of the high immersion VR treatments and the control condition (360_NEG vs. CONTROL and 360_POS vs. CONTROL), but the low immersion treatments are not significantly different than the control. 360° virtual reality combined with a negative framing significantly increased donations on the intensive margin compared to the classic unidirectional videos with negative framing (VID_NEG vs. 360_NEG). Although the average donations in the positive 360° virtual reality (360_POS) treatment (26,800IDR) appear to be much higher than those in the classic unidirectional (VID_POS) treatment (14,600IDR), these are not statistically different^5^. We do not observe any statistically significant differences between the positive and negative framing for either the low visual immersion (VID_NEG vs. VID_POS) treatments or the high visual immersion (360_NEG vs. 360_POS) treatments.

**Figure 5.**
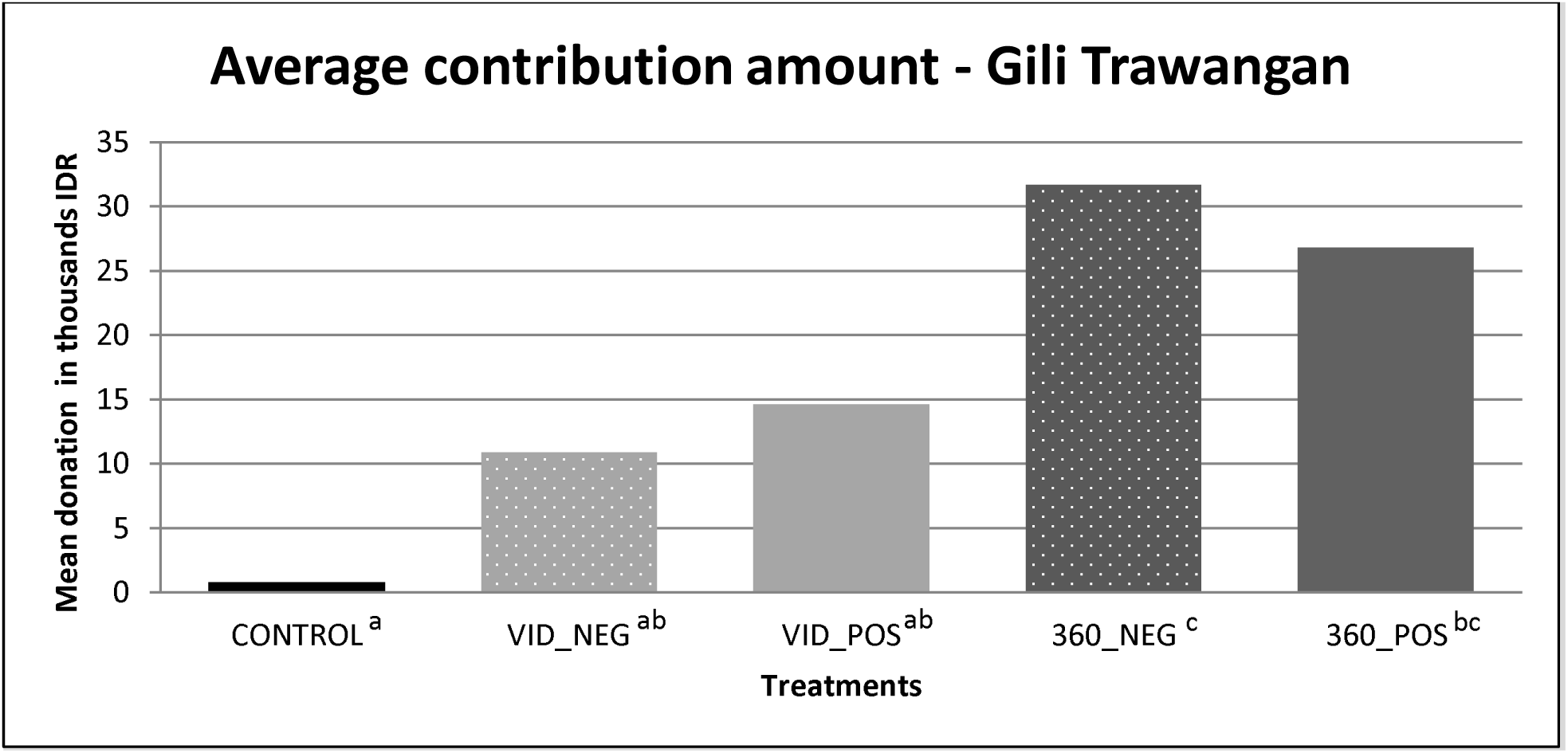
Average contribution amounts from tourists on Gili Trawangan. Treatment conditions with different superscript letters are significantly different at p<0.05; those with the same superscript letters are not statistically different from each other using the one-way ANOVA with Bonferroni for multiple comparisons.

When analyzing the donation amount as a dependent variable using an OLS regression, the data show that age and distance that a person lives from the sea in kilometers are both significant indicators of donation behavior. The older a person is and the further they live from the coast is correlated with higher donation amounts. Although just on the boarder of significance at p=0.052, a person’s self-perception of their level of environmental consciousness was correlated with donation amount. Interestingly, there is a negative relationship between environmental consciousness and donation amount, but this result is not significant at p<0.05. Confirming the results of the treatment comparison, we see significant differences between the highly immersive VR treatments (360_NEG and 360_POS) compared to the omitted CONTROL variable. Surprisingly there is not a significant effect of the level of income on donation amount as was seen in the first study. Unlike the Bogor sample, we do not see a significant effect of the level of income on the donation amount.

### 4.3 Emotional Responses: Pooled sample

The questions in the survey eliciting emotional responses and sense of presence were provided with a Likert scale presenting seven ordered response levels with labeled poles. The positive poles were coded with 1, and the negative poles with 7, respectively. We compare the mean scores for negative (NEG) versus positive (POS) valence and low immersive virtual experience (VID) versus high immersive virtual experience (360). We have pooled the data across the Bogor and Gili Trawangan samples for this analysis since we used the same treatments and the same survey questionnaires in both sample populations. Referring to Table 7, there are statistically significant differences showing that people in the positive treatment feel less sad compared to those in the negative treatment. People in the positive treatment and those in the VR treatment (360) reported feeling more happy than those in the negative treatment and those in the classic video treatment. Respondents in the negative treatments and in the VR treatments felt on average more helpless, but also more hopeful compared to those in the positive framing and video treatments. Participants exposed to the 360° virtual reality film reported feeling significantly more worried about global climate change than their counterparts in the classic video treatments.

**Table 5.** Linear regression of Bogor data with the donation amount as the dependent variable.

| Independent Variables | Donation amount | Donation amount | Donation amount |
| --- | --- | --- | --- |
| Age | **0.687 (3.53)\*\*** | **0.663 (3.42)\*\*** | **0.463 (2.52)\*** |
| Female | 0.997 (0.41) | 2.316 (0.94) | 3.974 (1.71) |
| Education | 0.215 (0.19) | -0.467 (0.41) | 0.377 (0.35) |
| KM to sea | 0.003 (1.81) | 0.003 (1.62) | 0.003 (1.53) |
| Income | **1.735 (2.96)\*\*** | **1.559 (2.64)\*\*** | **1.733 (3.11)\*\*** |
| Religiosity | 1.320 (1.27) | 0.273 (0.24) | 0.297 (0.28) |
| Env index |  | 1.555 (1.03) | 1.876 (1.32) |
| Optimist |  | -0.057 (0.06) | 0.243 (0.28) |
| Seen reefs |  | 1.691 (0.89) | 1.682 (0.94) |
| Env consciousness |  | 1.500 (1.35) | 0.946 (0.90) |
| Generosity |  | 1.030 (0.90) | 0.817 (0.75) |
| Sustainable future |  | **2.287 (2.01)\*** | 1.833 (1.71) |
| Control |  |  | 0.000 |
| VID_NEG |  |  | **23.334 (6.55)\*\*\*** |
| VID_POS |  |  | **23.340 (6.42)\*\*\*** |
| 360_NEG |  |  | **27.422 (7.65)\*\*\*** |
| 360_POS |  |  | **28.542 (7.94)\*\*\*** |
| Constant | 41.12 (4.81)** | 18.22 (1.49) | 1.16 (0.10) |
| $R^2$ | 0.04 | 0.06 | 0.18 |
| $N$ | 642 | 627 | 627 |

$t$ statistics in parentheses; \* $p < 0.05$ ; \*\* $p < 0.01$ ; \*\*\* $p < 0.001$
Note: Indicators obtained from questionnaire. Variables: Age (continuous), Female (Dummy=1 if Female), Education (scale of 1 elementary school to 6 post-graduate education) KM to sea (continuous), Income (perception scale of 1 lowest to 7 highest income), Religiosity (self-identified on a scale of 1 not at all to 7 most religious), Env Index (mean based on scaled responses to 10 environmental behavior indicator variables), Optimist (perception regarding climate change on a scale of 1 optimistic to 7 pessimistic), Seen Reefs (Dummy=1 if have seen reefs in the ocean), Env consciousness (perception of environmental consciousness compared to other people of the same age on a scale of 1 lowest to 7 highest), Generosity (perception of generosity compared to other people of the same age on a scale of 1 lowest to 7 highest), Sustainable future (perception of living a sustainable and environmentally conscious lifestyle on a scale of 1 not at all sustainable to 7 completely sustainable), Control (Dummy=1 in control treatment) is omitted treatment dummy, Negative (Dummy=1 if in treatment Unidirectional Negative message film), Positive (Dummy=1 if in Unidirectional Positive message film), 360 Negative (Dummy=1 if in 360 degree Negative message film, 360 Positive (Dummy=1 if in 360 degree Positive message film).

**Table 6.** Linear regression of Gili Trawangan data with donation amount as the dependent variable.

| Independent Variables | Donation amount | Donation amount | Donation amount |
| --- | --- | --- | --- |
| Age | **0.883 (3.90)\*\*** | **1.031 (4.30)\*\*** | **0.900 (3.78)\*\*** |
| Female | 0.105 (0.02) | -0.100 (0.02) | 0.687 (0.14) |
| Education | -0.947 (0.44) | -2.466 (1.09) | -3.033 (1.35) |
| KM to sea | **0.006 (2.43)\*** | **0.007 (2.58)\*** | **0.006 (2.36)\*** |
| Income | 1.479 (0.90) | 0.359 (0.20) | 0.697 (0.39) |
| Religiousness | -1.029 (0.84) | -1.020 (0.75) | -1.084 (0.82) |
| Env index |  | 0.212 (0.06) | 0.673 (0.20) |
| Optimist |  | 1.768 (1.18) | 0.844 (0.56) |
| Dive |  | 3.772 (1.02) | 3.257 (0.90) |
| Seen reefs |  | 5.467 (0.81) | 2.287 (0.35) |
| Env consciousness |  | -3.368 (1.78) | -3.614 (1.96) |
| Generosity |  | 3.902 (1.79) | 3.577 (1.67) |
| Sustainable future |  | 2.161 (1.09) | 0.613 (0.31) |
| Control |  |  | 0.000 |
| Negative |  |  | 9.508 (1.23) |
| Positive |  |  | 12.308 (1.61) |
| 360 Negative |  |  | **25.429 (3.28)\*\*** |
| 360 Positive |  |  | **21.944 (2.96)\*\*** |
| Constant | -8.916 (0.65) | -26.847 (1.12) | -21.239 (0.89) |
| $R^2$ | 0.10 | 0.16 | 0.23 |
| $N$ | 201 | 185 | 185 |

$t$ statistics in parentheses; \* $p < 0.05$ ; \*\* $p < 0.01$
Note: Indicators obtained from questionnaire same as Table 5 except Dive (Dummy=1 if scuba dive on Gili Trawangan).

**Table 7.** Table showing t-test results comparing negative framing (NEG) to positive framing (POS) and low visual immersion (VID) to high visual immersion (360)

|  | NEG vs POS |  | VID vs 360 |  |
| --- | --- | --- | --- | --- |
| Sad | 3.87 | 4.36 *** | 4.13 | 4.10 |
| Happy | 3.34 | 3.11 * | 3.35 | 3.10* |
| Helpless | 4.57 | 4.97* | 4.90 | 4.64* |
| Hopeful | 2.38 | 2.07*** | 2.40 | 2.07*** |
| Reflective | 1.82 | 1.74 | 1.80 | 1.76 |
| Worried | 1.38 | 1.41 | 1.45 | 1.34** |
| Part of the underwater world | 2.25 | 2.46* | 2.61 | 2.11*** |
| Surrounded by water | 2.16 | 2.21 | 2.51 | 1.87*** |
| Underwater | 2.59 | 2.50 | 3.00 | 2.1*** |
| Global consequences | 1.40 | 1.36 | 1.44 | 1.32** |
| Personal consequences | 2.01 | 1.92 | 2.03 | 1.90* |

\*p<0.05, \*\*p<0.01, \*\*\*p<0.001

As is expected, those exposed to the 360° virtual reality film felt a significantly higher sense of presence than those in the classic video treatment. Presence was measured by the questions ‘Did you feel like you were part of the underwater world?’; ‘Did you feel like you were surrounded by corals and fish?’; ‘Did it feel like you were really under water?’. Additionally, those that experienced the 360° virtual reality film perceived significantly higher global and personal consequences from global climate change.

When comparing the control condition to the negative and positive message valence and the low and high visual immersion, we see significant differences on almost every indicator of emotional feeling and sense of presence (see Table 8). The only indicators that do not appear to be significantly different from the control condition are those responses in the positive frame and 360° films to feeling hopeful.

**Table 8.** Table showing t-test results comparing the control to negative framing (NEG), positive framing (POS), low visual immersion (VID) and high visual immersion (360)

|  | Control | NEG | POS | VID | 360 |
| --- | --- | --- | --- | --- | --- |
| Sad | 5.28 | 3.87*** | 4.36*** | 4.13*** | 4.1*** |
| Happy | 2.53 | 3.34*** | 3.11*** | 3.35*** | 3.1*** |
| Helpless | 6.02 | 4.57*** | 4.97*** | 4.9*** | 4.64*** |
| Hopeful | 2.13 | 2.38** | 2.07 | 2.4** | 2.07 |
| Reflective | 2.73 | 1.82*** | 1.74*** | 1.8*** | 1.76*** |
| Worried | 2.13 | 1.38*** | 1.41*** | 1.45*** | 1.34*** |
| Global consequences | 1.67 | 1.4*** | 1.36*** | 1.44*** | 1.32*** |
| Personal consequences | 2.47 | 2.01*** | 1.92*** | 2.03*** | 1.9*** |

\*p<0.05, \*\*p<0.01, \*\*\*p<0.001

In all other cases, compared to the control, those in the treatment conditions felt overall significantly more sad, less happy, more helpless, less hopeful (only NEG and VID conditions), more reflective, more worried about climate change, and more strongly that climate change would have global and personal consequences.

## 5. Discussion and conclusion

In this paper we employ field experiments to examine the effects of virtual reality and message framing on conservation behavior and emotions. We surveyed two populations – an Indonesian sample in Bogor, Indonesia and a tourists sample in Gili Trawangan, Indonesia. We measure real behavior using donations to a conservation charity organization. We find that communicating an issue through the use of virtual reality technology is more effective at increasing donations than written donation requests, but the level of effectiveness is dependent on the target population and the immersive experience. The most effective treatment for the tourist population in Gili Trawangan was the 360° VR film with a *negative* audio message. This treatment increased donations by more than fifty percent compared to the classic video treatments. Although there was a fifteen percent increase in donations from the positive message framing, this difference was not statistically significant. The tourists differentiated their behavior between the type of media they experienced. The number of people donating was significantly higher in the classic video treatments compared to the control. And this effect increased significantly when participants were exposed to the 360° virtual reality video compared to those that saw the classic video. In our Bogor sample, however, the 360° VR film with *positive* message framing garnered the highest average donation amounts by at least a five percent margin.

In Bogor, all forms of media (classic unidirectional video and 360° VR video) led to significant increases in the average contribution amounts compared to no media communication (control). This result differed in the Gili Trawangan population where respondents in all treatments gave more, but only those in the 360° VR treatments gave significantly more on average than those in the control. We try to explain this difference given that we used the VR head mounted displays to screen all video treatments and the personal familiarity with the technology and the experience created with the technology may be different between the populations. The VR head mounted display creates a relatively immersive environment even without the video’s 360° responsive motion capability. It does so by blocking out other visual and auditory distractions and through physical proximity. The Bogor sample may have been more sensitive to the overall immersion created by the VR head mounted display due to the fact that this is a relatively new piece of technology that not many people in Bogor may have experienced before. In fact, we observed that many people froze their gaze locked in to looking in one direction only during the 360° VR treatments, which essentially limits a 360° experience to the experience provided by the classic video treatment. The nature of the technology physically displays the content of the video proximally very close to an individual within just a mere few centimeters away from a person’s eyes. Far fewer participants in the Bogor sample have seen a coral reef in real life compared to the sample in Gili Trawangan (Bogor = 51% and Gili Trawangan = 83%). The realistic and very close up underwater experience may have been the first time some have seen a glimpse into life underwater. Upon donning the headset, one participant in Bogor actually shrieked and was scared because she didn’t know how to swim and felt physically and emotionally like she was in danger because she felt as if she was in the water. We also had many comments from divers in Gili Trawangan that the experience really felt like they were diving and we even saw some attempting to equalize the pressure in the ears as the virtual descent underwater begins. So an increase in familiarity with the VR headset technology and more familiarity with underwater experiences, may explain the differences we see between the two samples. The tourists from Gili Trawangan appear to be more sensitive to the differences between viewing a classic video using a VR headset and viewing a 360° video with a VR headset. This is reflected in the significant differences observed in donations on the extensive and intensive margins in the Gili Trawangan sample but not in the Bogor sample.

We do not find significant differences in donation behavior between positive and negative message framing when paired with visual immersion (i.e., VID_POS vs. VID_NEG or 360_POS vs. 360_NEG) in either of the sample populations. The differences in the framing were purposefully subtle focusing only on a portion of the script towards the end of the video. The majority of research focusing on message framing is limited to text and still visual images (O’Keefe and Jensen 2007) with very few exceptions examining video messages (Rothman et al. 2006, Apanovitch, McCarthy, and Salovey 2003, Schneider et al. 2001). Therefore, relatively little is known about how people process positively and negatively framed information in the context of moving images, and especially in the context of VR. A negative relationship between virtual reality and memory recall has been documented in other studies which was attributed to underlying mechanisms such as limited cognitive capacity and mediated arousal (Bailey et al. 2012). Advancing from written text alone to text with still images to moving images with audio already represents a considerable increase on one’s cognitive burden to process information. Adding yet another layer of immersion through the VR headset and increased field of vision to 360° may have resulted in less capacity to process the audio information thereby lowering any effect of framing. The visual attention grabbing capabilities of immersive visual experiences may drain mental resources related to audio attention and mental processing. Human brains have limited cognitive resources to navigate through the world, including perceiving virtual experiences. According to a study by Lang (2006) using the Limited Capacity Model of Motivated Mediated Message Processing, our brains are continuously allocating processing resources to encode, store, and retrieve information during mediated message use as a function of the structure, content, motivation, and personal relevance of the message. Put simply, media experiences that may strain cognitive resources (i.e., underwater scenes represent highly vivid and unusual imagery with constant movement in an alien environment) could mean that people hear the audio message but are unable to process the content of the information and what they should do about it. Based on participants’ responses to the emotional and presence questions, it is evident that the framing has an effect on their feelings, it just doesn’t appear to lead to behavioral action in the form of increased donation amounts. Perhaps pairing the audio narration with written subtitles to reinforce the salience of the message would lead to behavior change.

Both positive and negative framing, and low and high visual immersion significantly affect emotional responses in comparison to no media communication. In Bogor, the video in all cases (VID and 360) heightened the awareness and concern about climate change and fostered higher donations. The tourist population in Gili Trawangan also reported increased emotional concern in all the treatment cases compared to the control. Increased emotions, however, did not necessarily translate symmetrically to action across the treatments. In all cases, those in the 360° treatments differed significantly in their donation behavior from the control. The framing had more of an effect on one’s emotional feeling but there was no difference between the positive and negative framing on how one perceived global climate change. In line with similarly framed experimental studies, we conclude that positive and negative frames can have systematic impacts upon both emotional responses and actual behavior (Rothman et al. 2006). However, there appear to be confounding factors that determine how framing plays a role and these are not well understood in a climate change context. Our study found mostly no differences in the framing except in one’s reported emotional state. It should be noted that studies that focus only on measuring emotion or intention as a proxy for action, may be overstating the behavioral impact of framing interventions and without additional content, changes in framing alone may not be effective at behavior change. And worse, focusing only on a particular type of framing ignores the nuances in the decision making process and between individuals that may require different types of messages targeted at different audiences to evoke emotion and change behavior. There are rarely ‘one-size fits all’ approaches to communicating complex issues such as climate change and messages should be tailored to fit the needs of the organization and the target audience. As noted by Lorenzoni, Nicholson-Cole, and Whitmarsh (2007), considering climate change risk involves judgments about uncertain and complex science, potential future impacts, and personal perceptions and values. Therefore, some context is always necessary when presenting information about climate change (Hulme 2008), but a positive or negative message valence does not seem to make much of a difference in our study. At least in the case of tourists, we can say that the negative framing is most effective when combined with virtual reality technology.

The results of the present study hold a complex set of implications for climate communication – but ones which we believe are broadly in line with the existing wider literature on framing and virtual reality communication. Communicating the crisis of coral reefs using visual information is related to an increased sense of presence and emotion, with the highest scores in the 360° VR treatment. This study provides empirical support that the use of 360° video using VR head-mounted devices can have the potential to attract more people to donate, and increase donations. When trying to communicate important video messages while fundraising in public where there can be many distractions, our research would also support the use of immersive portable viewing devices such as VR headsets even with the use of classic unidirectional videos. The VR devices may attract people and the immersive and individual experience created by the device may be sufficient to drive donations even without the increased immersion created by a 360° viewing experience. Further, our study shows that some observations, but not all, from VR behavioral studies in the lab translate to behavior in the field and the same is true with lessons learned from message framing studies taken from other domains when translated into the context of climate change. Given the relative infancy of this realm of literature, it is necessary to continue to conduct research examining the effects of virtual reality technology on human behavior, as well as the impacts of positive and negative message valence in the context of climate change. This is especially important during a time when information sources on these subjects are dominated by popular media and anecdotal evidence as opposed to evidence-based research.

## Acknowledgements

First and foremost, we want to thank Claudia and Hendrik Schmitt from “The Jetlagged” for the outstanding underwater film production. The research was co-funded by the Germany Federal Ministry of Education and Research (BMBF) university competition for the “Sea and Ocean” science year 2016-2017 and the Leibniz Association SAW-2014-ZMT-1 317 “Triple C: Contributions to Coral Commons” Project. We are grateful to the Institut Pertanian Bogor (IPB) and Dr. Luky Adrianto for helping facilitate this research. This research would not have been possible without the dedicated and hard-working master’s students from IPB and interns from Gili Eco Trust that worked as research assistants throughout this process. Special thanks to Delphine Robbe and Siân Williams of the Gili Eco Trust who provided their expertise and supported our activities to make this research possible.

## Contribution statement

The following contributions were provided by the authors: Katherine Nelson conceived of the research idea, acquired funding from BMBF, designed the research, wrote the video script and conceived the video content, conducted the field research and data collection process, analyzed and interpreted the data, and wrote the manuscript. Eva Anggraini conducted the field research and data collection process. Achim Schlueter acquired Triple C funding through the Leibniz Association, designed and conducted the field research, and critically revised manuscript content.

## Footnotes

1 Herein the terminology “virtual reality” includes 360° films viewed using head-mounted displays which provide an immersive experience. Although there is some discussion disputing the use of the term “virtual reality” to describe 360° films viewed using VR headsets, the term is widely accepted among popular media, the general public, the consumer-electronic industry, marketing companies, and charity organizations.

2 Please visit the link to view the full 360° (negative valence) video of *Coral Reefs: Life Below the Surface* produced by ‘The Jetlagged’ in collaboration with the Leibniz Centre for Tropical Marine Research (ZMT) https://www.youtube.com/watch?v=2TPG8lcfeDc&t=64s

3 The paper was used to clean up a mess created as part of the experimental design.

4 Exchange rate on April 4, 2018 1USD=13762.24IDR or 1EUR=16902.1IDR. Source: https://www.exchange-rates.org/HistoricalRates/P/EUR/4-4-2018

5 The Bonferroni correction for multiple comparisons is known to be very conservative. When we relax the restrictions and compare across two treatments only using t-tests, significant differences are revealed between the VID_NEG vs. CONTROL (p<0.01), VID_POS vs. CONTROL (p<0.01), and between 360_POS vs. VID_POS (p<0.05).

