## Appendix for "Virtual reality as a tool for environmental conservation and fundraising"

360° video script

### "Coral Reefs: Life Below the Surface"

Written by Katherine Nelson and Claudia Schmitt
Directed by Hendrik Schmitt
Produced by Leibniz Centre for Tropical Marine Research (ZMT)

You are on the deck of the exploration vessel Shakti.

Together we are on a journey through the Coral Triangle, where the Indian Ocean and the Pacific meet.

You can't see it from the surface, but the Ocean teems with life.

Under our feet swim huge schools of fish, sea turtles and playful dolphins.

A dive on a coral reef is a voyage to another world.

The surrealistic landscape is shaded in blue and surrounded by life... Life in a thousand forms.

Come with us and dive into the amazing underwater world.

Just look around. Everywhere you turn, there are more coral reef fish here than anywhere else in the world!

Coral reefs are spectacular to behold, lush gardens in the sea, supporting a staggering amount of life in a densely packed, thriving marine metropolis.

In fact, coral reefs have the largest abundance and greatest diversity of life living together of any place on Earth, including the tropical rain forests.

The reef itself is a living, growing organism – a colony of tiny animals all working together to create the largest structures on Earth.

The Coral Triangle truly is the epicenter of marine biodiversity and also home to invertebrates, whales, sharks and 6 out 7 species of sea turtles.

As the planet keeps warming, our oceans are changing.

Coral bleaching, ocean acidification, overfishing and pollution are threatening the world's coral reefs and their fragile ecosystems.

Can you imagine an ocean without reefs and abundant sea life?

a) **"Positive" narration**
   Countless animal species, but also us humans depend on healthy coral reefs.

They provide food for millions, security and protection from storm surges, sustain livelihoods and host the potential for new medicines. They are essential for the balance in the oceans.

We must protect it before it's too late.

Each person and each action makes a difference. Together we can save the ocean for a bright and blue future of our planet. If we act now, we can save the coral reefs.

b) **"Negative" narration**
Countless animal species, but also us humans will suffer if coral reefs are destroyed.

Millions will lose their source of food and their livelihoods. Storm surges will hit the coastlines with full force. Potential cures for human diseases that might be found in the ocean could be lost forever.

If we don't protect the ocean now, it might already be too late.

If we don't change the way we treat our planet, pollution and plastic waste will accumulate. Carbon dioxide will proliferate, driving global warming to higher and higher levels. If we don't act now, coral reefs will likely be gone within the next 30 years.
